# Gcn2 rescues reprogramming in the absence of Hog1/p38 signaling in *C. neoformans* during thermal stress

**DOI:** 10.1101/2024.06.11.598457

**Authors:** David Goich, Amanda L. M. Bloom, Sean R. Duffy, Maritza N. Ventura, John C. Panepinto

## Abstract

The fungus *Cryptococcus neoformans* is an opportunistic pathogen of people that reprograms its translatome to facilitate adaptation and virulence within the host. We studied the role of Hog1/p38 in reprogramming translation during thermal stress adaptation, and found that this pathway acts on translation via crosstalk with the Gcn2 pathway, a well-studied regulator of general translation control. Using a combination of molecular assays and phenotypic analysis, we show that increased output from the Gcn2 pathway in a Hog1 deletion mutant is associated with rescue of thermal stress adaptation at both molecular and phenotypic scales. We characterize known outputs of the Hog1 pathway during thermal stress as either Gcn2-dependent or Gcn2-independent, and demonstrate that Hog1 activation regulates the Gcn2 pathway even in the absence of thermal stress. Finally, we implicate this phenomenon in another Hog1-regulated process, morphogenesis, and recapitulate Hog1-Gcn2 crosstalk in the distantly related fungal pathogen, *Candida albicans.* Our results point to an important link between the stress response machinery and translation control, and clarify the etiology of phenotypes associated with Hog1 deletion. More broadly, this study highlights complex interplay between core conserved signal transduction pathways and the utility of molecular assays to better understand how these pathways are connected.

**Importance:** *Cryptococcus neoformans* is an opportunistic pathogen of people that causes deadly cryptococcal meningitis, which is is responsible for an estimated 19% of AIDS-related mortality. When left untreated, cryptococcal meningitis is uniformly fatal, and in patients receiving the most effective antifungal regimens, mortality remains high. Thus, there is a critical need to identify additional targets that play a role in adaptation to the human host and virulence. This study explores the role of the stress response kinases Hog1 and Gcn2 in thermoadaptation, which is pre-requisite for virulence. Our results show that compensatory signaling occurs via the Gcn2 pathway when Hog1 is deleted, and that disruption of both pathways increases sensitivity to thermal stress. Importantly, our study highlights the insufficiency of using single gene deletion mutants to study gene function, since many phenotypes associated with Hog1 deletion were driven by Gcn2 signaling in this background, rather than loss of direct Hog1 activity.

## Introduction

*Cryptococcus neoformans* is a ubiquitous environmental yeast that causes deadly opportunistic infections in people who are immunocompromised. Cryptococcosis is responsible for an estimated 112,000 deaths annually, with a disproportionate impact in under-resourced areas such as Southern Africa [1]. Infection with *Cryptococcus* typically begins with primary infection of the lungs, followed by cryptococcal meningitis if pulmonary containment is unsuccessful [2]. If left untreated, cryptococcal meningitis is usually fatal. Even among individuals who receive the best available antifungal regimen, one-year mortality from cryptococcal meningitis remains at approximately twenty percent [1]. Thus, there is a critical need for discovery of novel approaches to mitigate mortality due to cryptococcosis.

The ability to tolerate mammalian endothermy is a pre-requisite for fungi to act as systemic human pathogens [3]. In addition to changes in transcription and metabolism, this process requires signaling to the translation apparatus and RNA decay machinery, reviewed in [4]. In response to many stressors, levels of basally abundant mRNAs such as ribosomal protein (RP) transcripts are repressed as part of the fungal environmental stress response [5]. This, coupled with transient repression of translation, frees ribosomes for reallocation to stress-related transcripts to drive efficient production of stress-mitigating proteins [6]. Defects in this process in fungal pathogens, called ***translatome reprogramming***, are associated with impaired stress adaptation and attenuated virulence [4]. We’ve documented reprogramming in response to several stressors that are relevant to infection in *C. neoformans*, and have identified key regulators of this process [7–10]. For example, *C. neoformans* lacking the major mRNA deadenylase, Ccr4, is unable to repress abundant growth-associated mRNAs and exhibits defects in survival at mammalian core temperature and attenuated virulence [9,11].

Previously, we identified two MAP kinase (MAPK) modules that are critical for remodeling of the cell wall and maintenance of β-1,3 glucan masking upon thermal stress [12]. Whereas the cell wall integrity MAPK module appeared to primarily regulate the transcriptional response to thermal stress, the high osmolarity glycerol (HOG) p38 MAPK pathway exhibited additional roles in post-transcriptional and translational regulation of the response [12]. Previous work on Hog1 homologs in model fungi suggests that the manner of regulation of stress responses varies substantially, even within the ascomycete lineage of fungi [13,14]. Thus, the mechanism of translatome regulation by Hog1 in basidiomycetes such as *C. neoformans*, which diverged from ascomycete model fungi approximately 400 million years ago [15], is especially unclear. While several studies of the Hog1 pathway in *C. neoformans* have identified the upstream machinery that regulates Hog1 activity [16–18], direct downstream targets of Hog1 in *C. neoformans* are not defined. Many studies including our own, however, have phenotypically associated Hog1-deletion with important processes related to stress response, virulence factor production, and translatome reprogramming [7,12,16,19].

In addition to the Hog1 pathway, we have also identified the kinase, Gcn2, as a key regulator of stress-adaptive translation in *C. neoformans* [8]. Gcn2 is canonically activated in response to ribosome collisions or accumulation of uncharged tRNAs caused by stress, resulting in global repression of translation initiation via phosphorylation of initiation factor 2α (eIF2α) [20]. This phosphorylation event globally inhibits cap-dependent translation initiation by limiting availability of the 43S pre-initiation complex (PIC), while paradoxically promoting increased translation of the coding sequence of the transcription factor Atf1/Gcn4, reviewed in [21]. The Atf1/Gcn4 transcriptional regulon, known as the integrated stress response (ISR), encodes proteins that are important for maintenance of amino acid biosynthesis, tRNA charging, and redox balance [22]. We recently characterized the role of Gcn2 and the ISR during stress in *C. neoformans*, and showed that this pathway is induced by mammalian core temperature [10].

Since the Hog1 and Gcn2 pathways appear to converge at the point of translatome reprogramming during thermal stress, the present study evaluates whether this is associated with crosstalk, compensation, or redundancy between these pathways. We found that levels of phosphorylated eIF2α are increased when the Hog1 pathway is disrupted, and that this is associated with increased translation repression in *hog1*Δ during thermal stress. We showed that Hog1 and Gcn2 both contribute to repression of abundant growth-associated mRNAs, which was defective when both pathways were disrupted in a double mutant. The double mutant also exhibited exacerbated defects in thermoadaptation, suggesting to us that this crosstalk is compensatory at both the molecular and phenotypic levels. Lastly, we show that crosstalk between these pathways can occur over a large evolutionary distance, and implicate compensation in driving other phenotypes of Hog1 deletion such as increased filamentation.

## Results

### Loss of Hog1 increases phosphorylated eIF2α during thermal stress in *C. neoformans*

Previous studies of ascomycete fungi suggest that Hog1/p38 is a regulator of global translational output in the context of several stressors, such as osmotic and oxidative stress [13,14]. In *C. neoformans*, we previously evaluated the role of Hog1 in the response to thermal stress at mammalian body temperature and found that the *hog1*Δ mutant exhibited more robust repression of global translation in response to thermal stress [12]. This somewhat unexpected result led to the hypothesis that an alternative translation regulation pathway is upregulated in the absence of Hog1.

Since our previous work has also established the Gcn2 pathway as a key regulator of translation in *C. neoformans* during stress [10], we began our studies by investigating whether eIF2α phosphorylation was increased in a Hog1 deletion mutant by western blotting (Fig. 1). We found that whereas the WT strain exhibited a modest level of eIF2α phosphorylation in response to thermal stress at mammalian core temperature, the *hog1*Δ mutant had robustly phosphorylated eIF2α (Fig. 1A). Increased basal eIF2α phosphorylation was also observed under unstressed conditions in *hog1*Δ, which were defined as mid-logarithmic growth at 30°C. The ratio of phosphorylated eIF2α signal to total eIF2α signal was greater in *hog1*Δ than WT (Fig. S1), indicating that a higher proportion of the protein is phosphorylated in *hog1*Δ.

**Fig 1:**
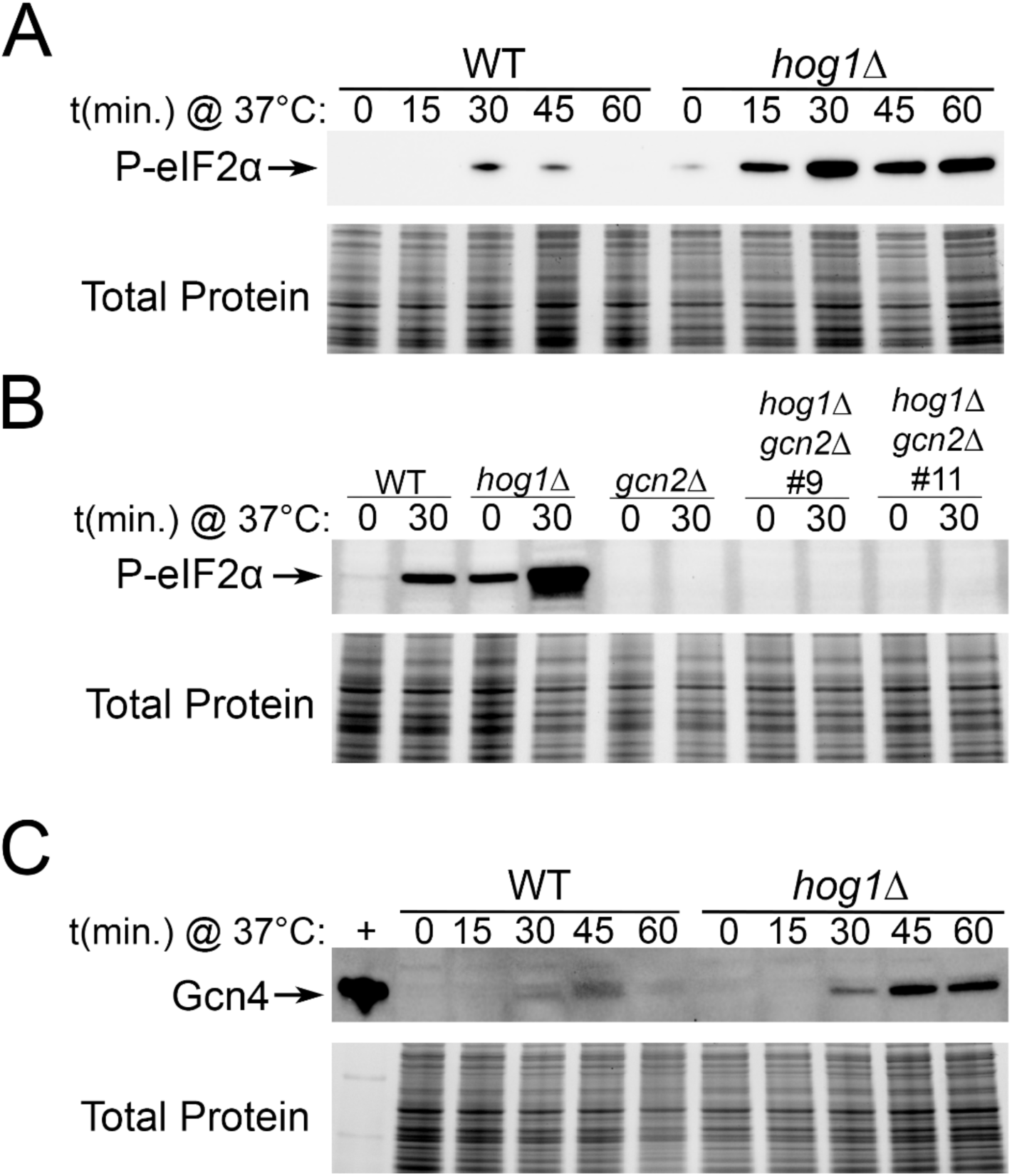
Loss of Hog1 increases Gcn2-dependent outputs during thermal stress. The indicated strains were grown to midlogarithmic phase at 30°C in YPD media, followed by resuspension in pre-warmed 37°C YPD media. Samples were collected at the indicated time points, and protein samples were prepared and probed by western blotting using an antibody against (A-B) phosphorylated eIF2α, or (C) Gcn4. Total protein is shown as a loading control. +; positive control recombinant Gcn4 protein. One representative replicate is shown (*n* = 3).

We previously described Gcn2 as the sole eIF2α kinase in *C. neoformans* [8]. To verify that Gcn2 alone was responsible for eIF2α phosphorylation even when Hog1 is deleted, we measured levels of phospho-eIF2α in WT, *hog1*Δ, *gcn2*Δ, and *hog1*Δ*gcn2*Δ mutants before and after thermal stress (Fig. 1B). Simultaneous loss of Hog1 and Gcn2 ablated eIF2α phosphorylation entirely, suggesting that upregulated eIF2α phosphorylation in *hog1*Δ is fully dependent on Gcn2.

Given the established role of the Gcn2 pathway in promoting translation of the integrated stress response (ISR) transcription factor, Gcn4, we next evaluated whether Gcn4 protein levels were increased in the absence of Hog1 (Fig. 1C). We found that the Hog1 deletion mutant had increased levels of Gcn4 during thermal stress, albeit not basally. Overall, these results demonstrate that outputs of Gcn2 activity, eIF2α phosphorylation and Gcn4 protein production, are increased in *hog1*Δ during thermal stress.

### Loss of both Hog1 and Gcn2 exacerbates defects in thermal stress adaptation

Since we saw evidence of increased Gcn2 signaling when the Hog1 pathway was disrupted, we next investigated whether this was associated with compensation at the phenotypic level. We hypothesized that increased eIF2α phosphorylation may serve a protective role in growth of the *hog1*Δ mutant during thermal stress.

To test this, we first employed spot plate analysis of WT, *hog1*Δ, *gcn2*Δ, and *hog1*Δ*gcn2*Δ under growth at ambient (30°C), mammalian core (37°C), and mammalian febrile (38°C) temperatures (Fig. 2A). We found that all mutants had reduced survival at mammalian febrile temperature compared to WT, with a more pronounced phenotype in the Hog1-deleted strains. The double-deletion mutant had few noticeable defects beyond the *hog1*Δ single mutant. This suggested that Gcn2 was not critical for survival of *hog1*Δ cells by spot plating.

**Fig 2:**
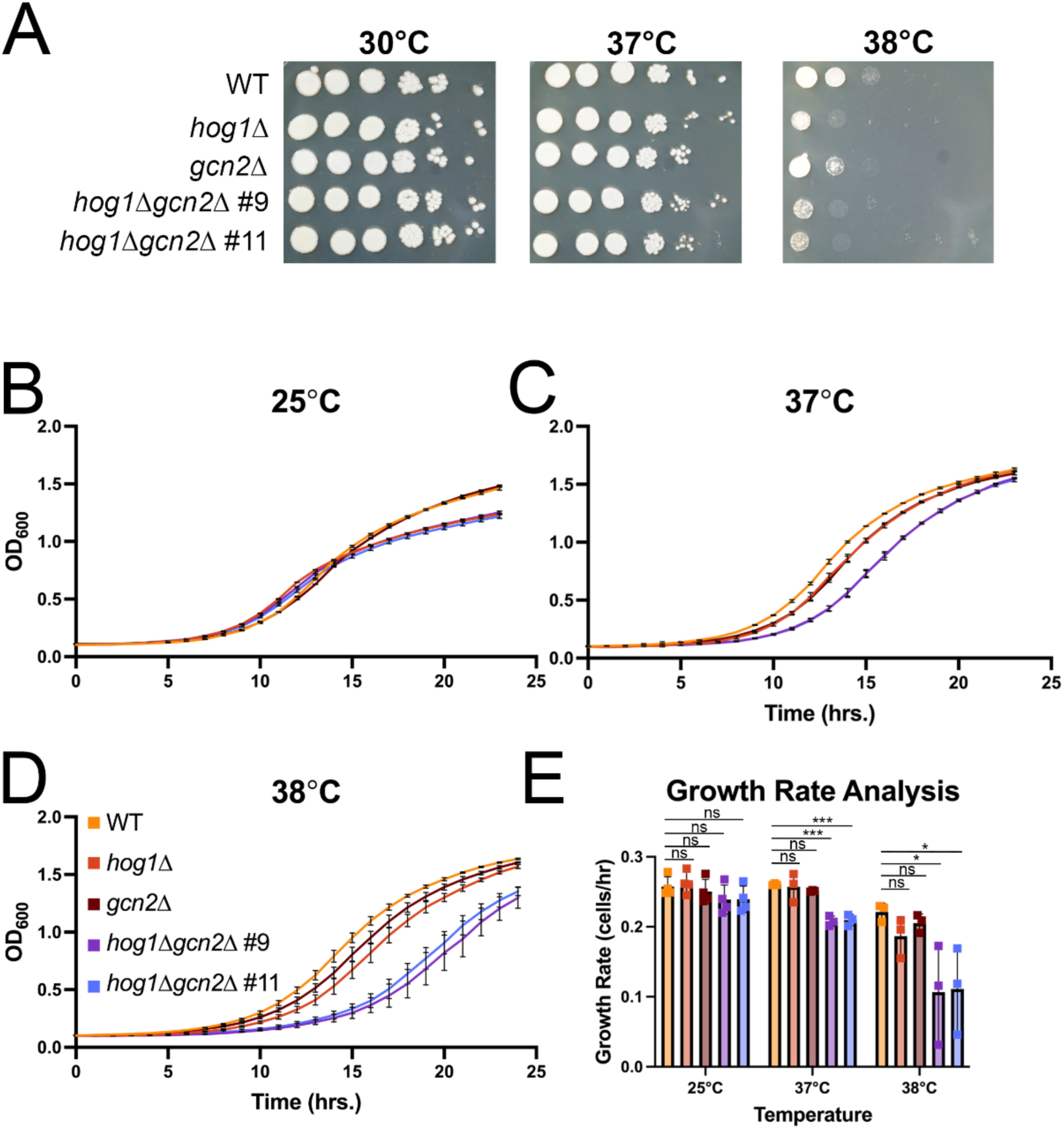
Simultaneous loss of Hog1 and Gcn2 exacerbates growth defects during thermal stress. The indicated strains were grown overnight in YPD media at 30°C. (A) Overnight cells were washed and resuspended in water to OD_600_ = 1.0. A ten-fold dilution series was performed, and 5μL of cells were spotted on YPD agar plates, incubated at the indicated temperatures, and photographed after 3 days. Shown is one representative replicate (*n* = 3). (B-D) Overnight cells were added to fresh YPD medium to an OD_600_ = 0.1 and 100μL of cells were plated in triplicate in a 96-well plate. Cells were grown with shaking at the indicated temperature, and the OD_600_ was read every hour. Each curve shows one representative biological replicate (*n* = 3), with error bars depicting standard deviation across triplicate wells. (E) The growth rate for the exponential portion of growth curves was calculated as described in the Materials and Methods. Error bars represent standard deviation across three biological replicates. Significance was calculated by one-way ANOVA with Dunnet’s test post hoc. *; p < 0.05; ***; p < 0.001.

In this experiment, however, we noted somewhat smaller colonies in the Hog1 deletion strains at elevated temperatures (Fig. 2A). This suggested to us that kinetic growth, rather than endpoint growth, may be of interest for study. Thus, we performed growth curve analyses at ambient (25°C), mammalian core (37°C), and mammalian febrile (38°C) temperatures (Fig. 2B-E). Whereas the *hog1*Δ, *gcn2*Δ, and *hog1*Δ*gcn2*Δ strains did not exhibit a substantial growth defect relative to WT at 25°C (Fig. 2B), elevating the temperature to mammalian core (Fig. 2C) and febrile (Fig. 2D) temperature increasingly stratified these strains. Overall, we saw relatively small temperature-dependent defects in growth of *hog1*Δ and *gcn2*Δ, which were exacerbated in the double knockout strains. The growth rate during exponential growth under each condition was calculated, and we observed a statistically significant difference in the growth rate of double knockout strains, relative to WT, at mammalian core and febrile temperatures (Fig. 2E). Collectively, our data show that both Hog1 and Gcn2 contribute to thermal stress adaptation, where loss of both uncovers a more severe kinetic growth defect at elevated temperatures.

### Cell wall and capsule dysregulation in *hog1*Δ is mostly independent of Gcn2

In our previous study we found that Hog1 was a regulator of cell wall reprogramming in the response to thermal stress [12]. To follow up on these findings, we investigated whether Gcn2 regulates cell wall remodeling in *hog1*Δ at elevated temperature.

We first evaluated the susceptibility of WT, *hog1*Δ, *gcn2*Δ, and *hog1*Δ*gcn2*Δ mutants to cell wall stressors by spot plate analysis at both ambient and mammalian core temperatures (Fig. 3A). Consistent with our previous findings [12], the *hog1*Δ mutant was sensitive to both caffeine and the detergent SDS. The *gcn2*Δ strain did not exhibit sensitivity to any of the cell wall perturbing agents tested. We found that the *hog1*Δ*gcn2*Δ strains largely phenocopied *hog1*Δ. Overall, this data suggests Gcn2 does not play a major role in the sensitivity of *hog1*Δ to cell wall stressors.

**Fig 3:**
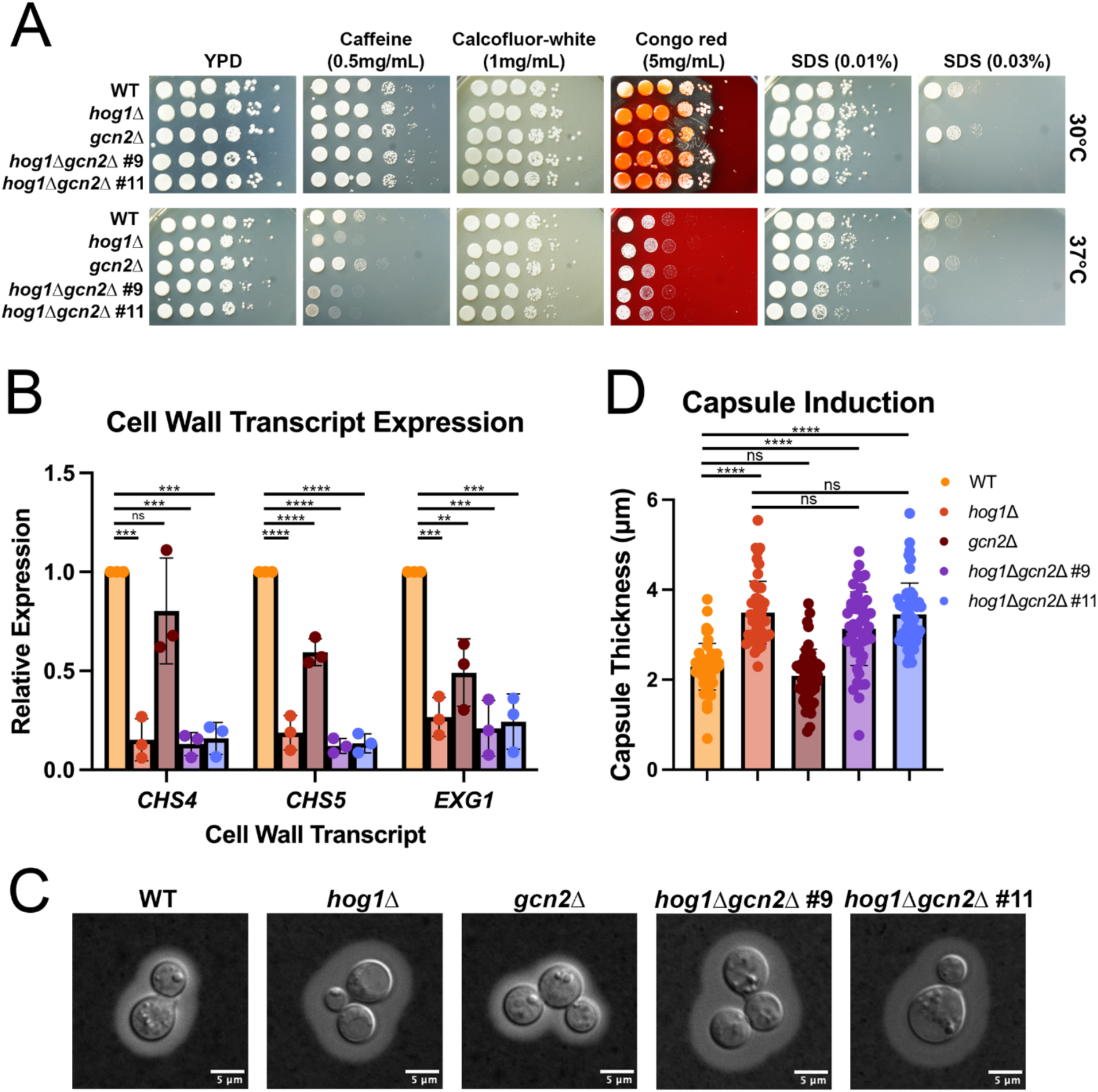
Gcn2 does not drive cell wall and capsule phenotypes associated with Hog1 deletion. (A) Cells grown at overnight at 30°C in YPD media were resuspended in water to an OD_600_ = 1.0. A ten-fold dilution series was performed, and 5μL of cells were spotted on YPD agar plates containing the indicated cell wall perturbing agents and incubated at the indicated temperatures for 3 days. No drug YPD plates serve as controls. One representative replicate is shown (*n* = 3). (B) The indicated strains were grown to midlogarithmic phase in YPD media at 30°C, followed by resuspension in 37°C YPD for one hour. RNA was extracted, followed by DNase I treatment, reverse transcription, and qPCR for the indicated transcripts. Differential expression was calculated by the ΔΔC_T_ method and normalized to the control transcript mitofusin. Significance was calculated by one-way ANOVA with Tukey’s test *post hoc* to correct for multiple comparisons. Error bars represent standard deviation across biological replicates (*n* = 3). **; p < 0.01, ***; p < 0.001; ****; p < 0.0001. (C) Representative DIC images of WT, *hog1*Δ, *gcn2*Δ, and *hog1*Δ*gcn2*Δ strains stained with India Ink after growth under capsule-inducing conditions. (D) Quantification of capsule size in WT, *hog1*Δ, *gcn2*Δ, and *hog1*Δ*gcn2*Δ strains (*n=50*). Significance was calculated by one-way ANOVA with Dunnet’s test post hoc. ****; p < 0.0001

Since several cell wall remodeling transcripts have lower expression in *hog1*Δ relative to WT during growth at 37°C [12], we next evaluated the expression of cell wall-related transcripts at mammalian core temperature by RT-qPCR (Fig. 3B). We measured levels of *CHS4* and *CHS5*, which encode chitin synthases, and *EXG1*, an exoglucanase, all of which are upregulated in response to thermal stress. Although loss of Gcn2 alone altered expression of some transcripts relative to WT, expression of these transcripts in *hog1*Δ*gcn2*Δ was not substantially different from *hog1*Δ. This suggests that Gcn2 regulates cell wall transcript expression in the WT background, but is not a major contributor to the expression levels observed in *hog1*Δ. Thus, this data does not support a compensatory role for Gcn2 in cell wall transcript expression when Hog1 is deleted.

Hog1 is also known to regulate capsule induction in *C. neoformans*, which is considered to be a virulence-defining trait for this pathogen. A previous study found that under capsule-inducing conditions, *hog1*Δ produces a significantly larger capsule than WT [16]. To determine whether altered capsule production in *hog1*Δ is attributable to the Gcn2 pathway, we induced capsule in WT, *hog1*Δ, *gcn2*Δ, and *hog1*Δ*gcn2*Δ and performed microscopy after India ink staining (Fig. 3C-D). We found that *hog1*Δ*gcn2*Δ was hypercapsular, and did not differ substantially from the *hog1*Δ single mutant. Additionally, we did not see a significant difference in capsule production in *gcn2*Δ compared to WT. This indicates that Gcn2 does not contribute to capsule production, whether or not Hog1 is present. Collectively, our data suggest that Gcn2 pathway signaling is not a major contributor to the cell wall or capsule phenotypes of *hog1*Δ.

### Gcn2 drives an altered translational response to thermal stress in *hog1*Δ

Since the Gcn2 pathway is a major regulator of translation initiation, we next began to investigate the role of crosstalk in regulating the translational response of *hog1*Δ to thermal stress. We previously showed that a *hog1*Δ mutant undergoes enhanced translation repression in response to thermal stress [12]. Given the increase in phospho-eIF2α in *hog1*Δ (Fig. 1), we hypothesized that enhanced repression in *hog1*Δ was a Gcn2-dependent phenomenon.

To test this, we performed polysome profiling in WT, *hog1*Δ, *gcn2*Δ, and *hog1*Δ*gcn2*Δ strains in response to thermal stress at 37°C (Fig. 4). Consistent with our previous studies [9,10,12], we observed translation repression in the WT upon thermal stress, with an increase in the free 60S ribosomal subunits and repression in the polysomes (Fig. 4A). By comparison, the response of *hog1*Δ was more robust, with greater repression throughout the polysome fraction (Fig. 4B). In the *gcn2*Δ mutant, accumulation of free 60S subunits was mostly ablated, with increased persistence of high molecular weight polysome complexes after stress (Fig. 4C). Profiles for the *hog1*Δ*gcn2*Δ strains appeared similar to the *gcn2*Δ single mutant (Fig. 4D). Importantly, the double knockout did not exhibit increased translation repression in response to thermal stress, as was seen in the *hog1*Δ single knockout. This suggested that the hyper-repressive response seen as a result of Hog1 deletion is dependent on the Gcn2 pathway.

**Fig 4:**
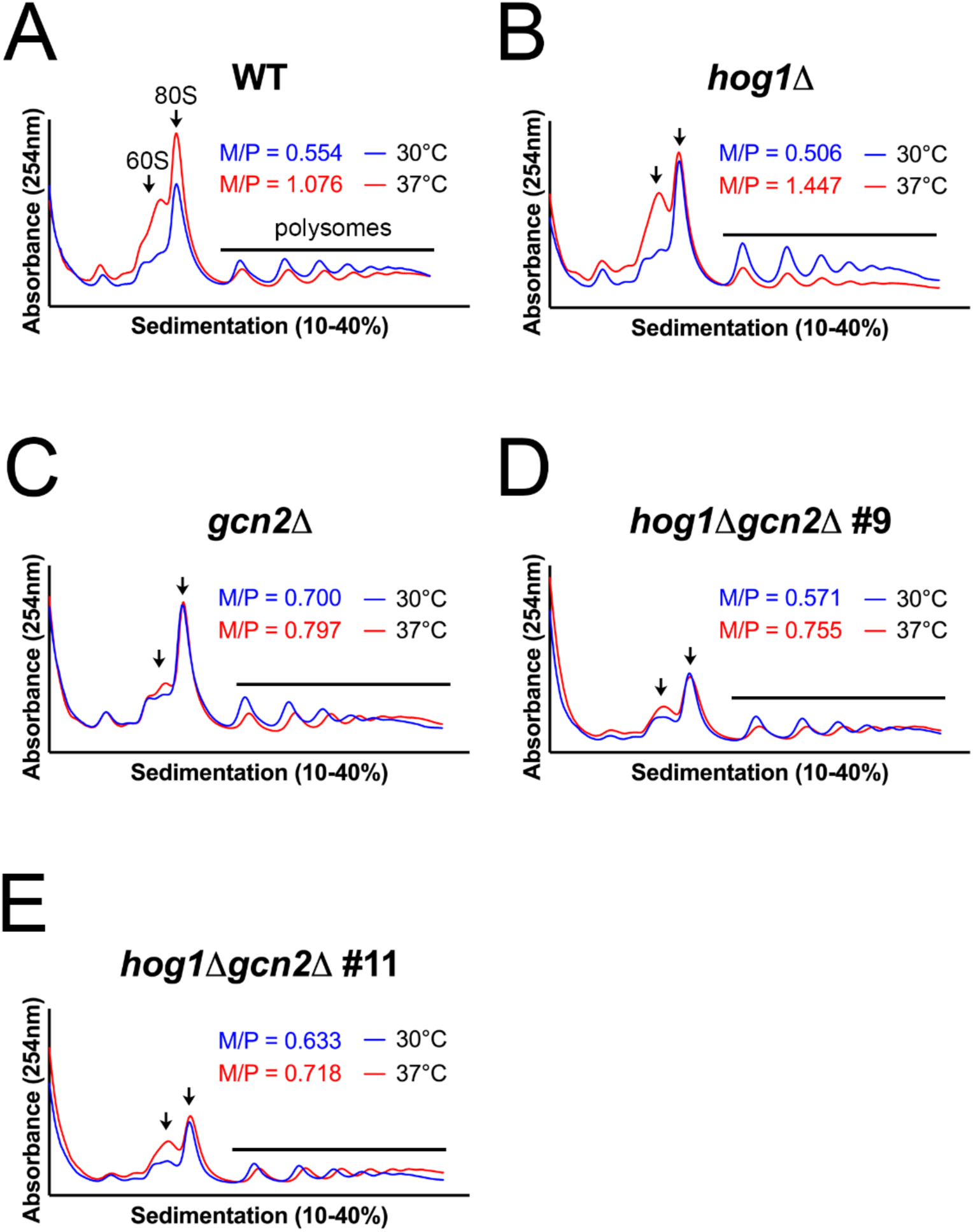
The altered translational response to thermal stress in *hog1*Δ is Gcn2-dependent. The indicated strains were grown to midlogarithmic phase in 30°C YPD media, then resuspended in 37°C media for thirty minutes. Cells were lysed the presence of cycloheximide and equivalent lysate (quantified by OD_280_) was centrifuged over a 10-40% sucrose gradient. Ribosomes and subunits were detected by measuring absorbance at 254nm throughout the gradient. Arrows point to peaks corresponding to (left) free 60S subunit and (right) mRNA-associated ribosome (80S, right). The black line covers the portion of the graph representing polysomes. The monosome-to-polysome (M/P) ratio was calculated as described in the Materials and Methods. Each graph depicts a representative biological replicate (*n* = 3).

The areas under the monosome and polysome portions of the profile were calculated, and the monosome-to-polysome (M/P) ratio was calculated for each curve as a measure of the extent of translation repression (Fig. S2). Although M/P quantification did not meet statistical thresholds for significance, the overall trend in the data agrees with our previous conclusions. The *hog1*Δ mutant exhibited a larger M/P ratio than WT upon thermal stress, which was ablated in the *hog1*Δ*gcn2*Δ mutant. Collectively, these data indicate that increased translation repression in *hog1*Δ is Gcn2-dependent.

### Hog1 and Gcn2 both contribute to repression of ribosomal biogenesis transcripts

Another key step in translatome reprogramming involves removal of abundant growth-associated transcripts from the translating pool of mRNAs. Bulk decay of these mRNAs, such as ribosomal protein (RP) transcripts, is necessary for efficient translation of the stress response [6]. Our previous work found that Hog1 pathway mutants exhibit prolonged repression of the RP transcript, *RPL2,* during thermal stress adaptation [12]. We hypothesized, again, that this enhanced response was dependent on the Gcn2 pathway.

To investigate whether the Gcn2 pathway contributes to RP transcript repression in *hog1*Δ, we performed northern blotting for the *RPL2* transcript during thermal stress in WT, *hog1*Δ, *gcn2*Δ, and *hog1*Δ*gcn2*Δ (Fig. 5A-D). The WT exhibited repression of *RPL2* over the first hour, followed by rebound over the second hour (Fig. 5A). The *hog1*Δ and *gcn2*Δ mutants were both able to repress *RPL2* to the same extent as WT after one hour, although recovery was somewhat defective (Fig. 5B-C). This was especially pronounced in *hog1*Δ, which maintained repression of *RPL2* even after two hours (Fig. 5C). Loss of both pathways, however, uncovered a defect in repression of *RPL2*, which was not repressed to the same extent as in WT (Fig. 5D).

**Fig 5:**
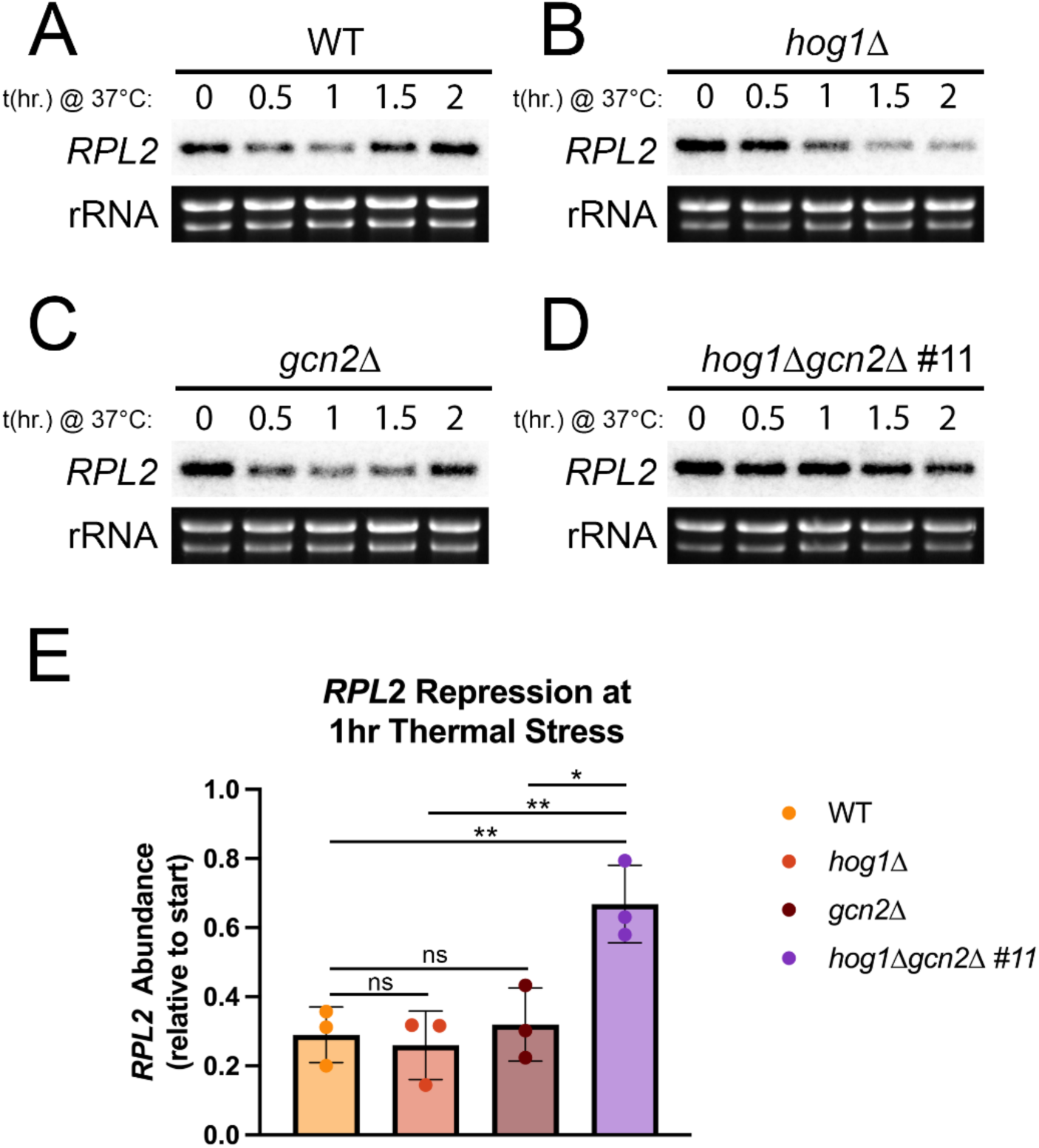
Hog1 and Gcn2 both contribute to repression of the abundant, growth-related transcript *RPL2*. (A-D) The indicated strains were grown to midlogarithmic phase in 30°C YPD media, then resuspended in 37°C YPD for two hours and time points were collected every thirty minutes. RNA was extracted and samples were probed by northern blot using a ^32^P-labeled oligonucleotide probe. rRNA is shown as a loading control. Shown is one biological replicate (*n* = 3). (E) *RPL2* signal was quantified by densitometry, normalized to rRNA, and the change after 1 hour was calculated. Error bars indicate the standard deviation across biological replicates (*n* = 3).

To quantify this difference, we measured the change in *RPL2* transcript abundance after one hour (Fig. 5E), which is the time of maximal repression in WT (Fig. 5A). There was a statistically significant defect in *RPL2* repression in the double knockout strain, compared to WT, *hog1*Δ, and *gcn2*Δ. While some degree of repression appeared to be independent of both pathways (Fig. 5D), these data indicate that both Hog1 and Gcn2 can regulate RP transcript abundance.

### Stimulation of the Hog1 pathway regulates levels of P-eIF2α

Since loss of Hog1 was associated with increased levels of phospho-eIF2α (Fig. 1), we next investigated whether stimulating the Hog1 pathway is sufficient to alter levels of phospho-eIF2α during stress. Since thermal stress acts on the Hog1 and Gcn2 pathways simultaneously [10,12], this stressor was not useful for testing cause-and-effect relationships between these pathways. To overcome this, we used the drug fludioxonil, which stimulates the Hog1 pathway by acting on its upstream regulators [23], and 3-amino-1,2,4-triazole (3-AT), which inhibits histidine biosynthesis resulting in Gcn2 activation.

To test whether Hog1 activity regulates levels of phospho-eIF2α, we treated WT cells with 3-AT to activate Gcn2, with or without addition of fludioxonil to activate Hog1, and performed western blot analysis (Fig. 6A-B). We found that simultaneous stimulation of the Hog1 pathway was associated with lower levels phospho-eIF2α in response to 3-AT (Fig. 6A), as well as lower levels of the downstream transcription factor Gcn4 (Fig. 6B). This data suggests that activation of Hog1 by fludioxonil reduces Gcn2-dependent outputs in the response to 3-AT.

**Fig 6:**
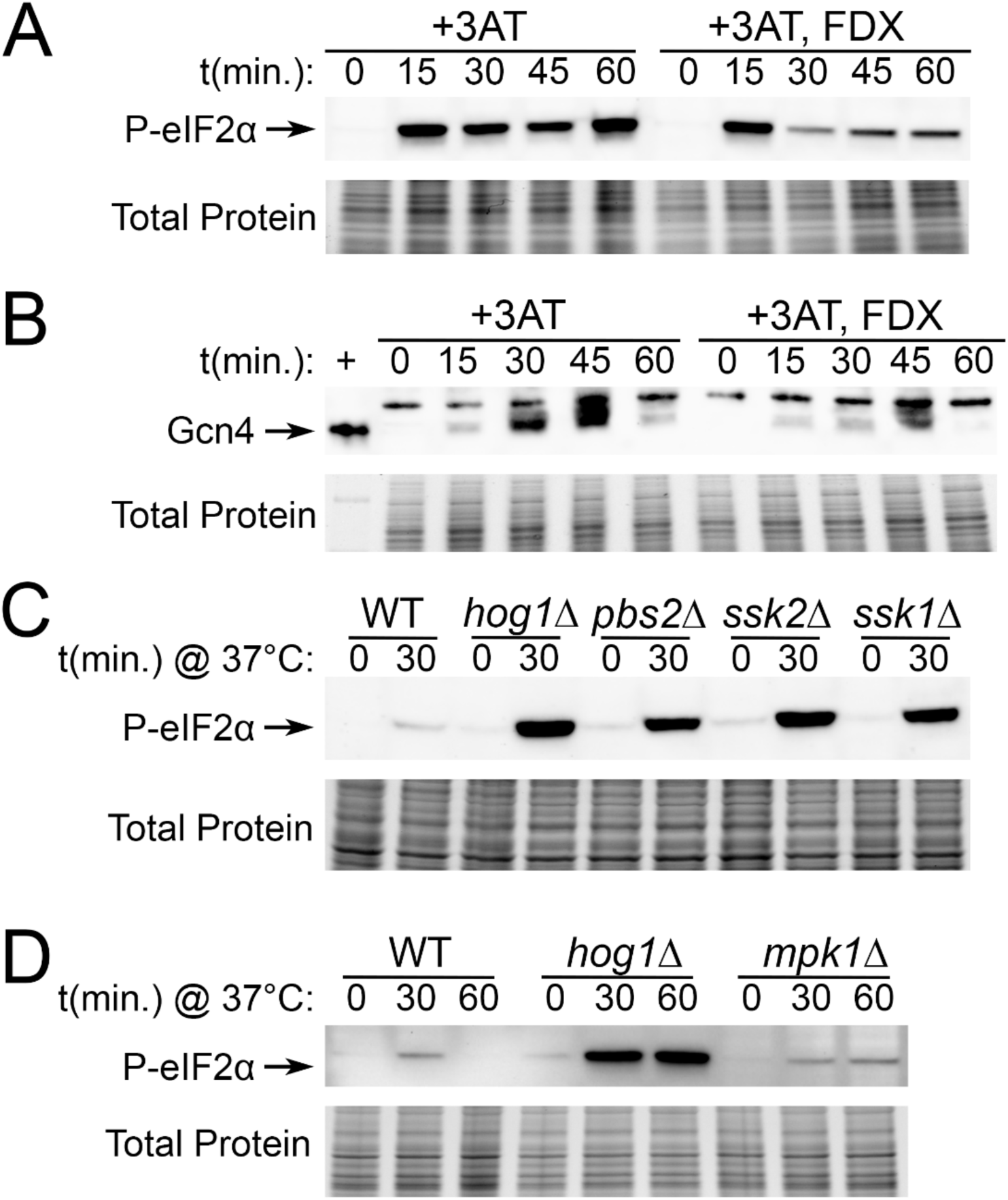
Stimulation of the Hog1 pathway regulates levels of P-eIF2α. (A-B) WT cells were grown to midlogarithmic phase in 30°C YPD media, then treated with 40mM 3-AT and 5μg/mL fludioxonil (FDX) as indicated. Time points were collected and protein lysates were probed for (A) phosphorylated eIF2α, or (B) Gcn4 by western blot. (C-D) The indicated strains were grown to midlogarithmic phase in 30°C YPD media, then resuspended in 37°C YPD for the indicated amounts of time. Protein lysates were probed for phosphorylated eIF2α by western blot. Total protein is shown as a loading control. +; positive control recombinant Gcn4 (*n = 3*).

We next set out to investigate whether upregulation of Gcn2-eIF2α signaling in *hog1*Δ during thermal stress is dependent on known upstream regulators of the Hog1 pathway. This was a relevant question to ask since the mechanism by which Hog1 participates in the thermal stress response is not well-characterized in *C. neoformans*. We found that deletion mutants of upstream members of the MAPK module, Pbs2, Ssk2, as well as the upstream two-component response regulator, Ssk1, all exhibited increased levels of phospho-eIF2α in response to thermal stress (Fig. 6C). This result indicates that both that crosstalk with the Gcn2 pathway during thermal stress is driven by canonical activity of the Hog1 pathway.

Lastly, having established a relationship between Hog1 pathway signaling and Gcn2 activity, we were curious whether this is a general feature of MAPK signaling. Our previous work demonstrated that the cell wall integrity MAPK, Mpk1, is also activated by thermal stress. Thus, we measured levels of phospho-eIF2α in response to thermal stress in an *mpk1*Δ mutant (Fig. 6D). The response of *mpk1*Δ, albeit prolonged, was similar in magnitude to WT and substantially less robust than in *hog1*Δ. This suggests that the Hog1 pathway is unique with regard to Gcn2 signaling. Together, these data show that crosstalk with Gcn2 is driven by the upstream components of the HOG pathway, and that stimulation of this pathway is sufficient to alter Gcn2-dependent signaling even in the absence of thermal stress.

### Hog1-Gcn2 crosstalk regulates morphogenesis and is conserved in *C. albicans*

Given the potential for Hog1 to regulate Gcn2 in contexts other than thermal stress, we next wanted to evaluate how this crosstalk may impact other Hog1-dependent processes. Since there is a well-established link between disruption of the Hog1 pathway and altered filamentation [16,24,25], we investigated whether morphology of *hog1*Δ is dependent on Gcn2. Although *C. neoformans* grows almost exclusively as a yeast, filamentation occurs during mating, and is considerably increased in the absence of Hog1 [16]. We mated *MAT*α WT, *hog1*Δ, *gcn2*Δ, and *hog1*Δ*gcn2*Δ with a *MAT*a strain (KN99a) and observed the extent of filamentation by microscopy after three and five days (Fig. 7A). As expected, the *hog1*Δ strain was hyperfilamentous, and produced substantially more filaments than WT. By contrast, the *gcn2*Δ and *hog1*Δ*gcn2*Δ strains filamented at comparable or lower levels than the WT. This indicates that hyperfilamentation of *hog1*Δ is dependent on Gcn2.

**Fig 7:**
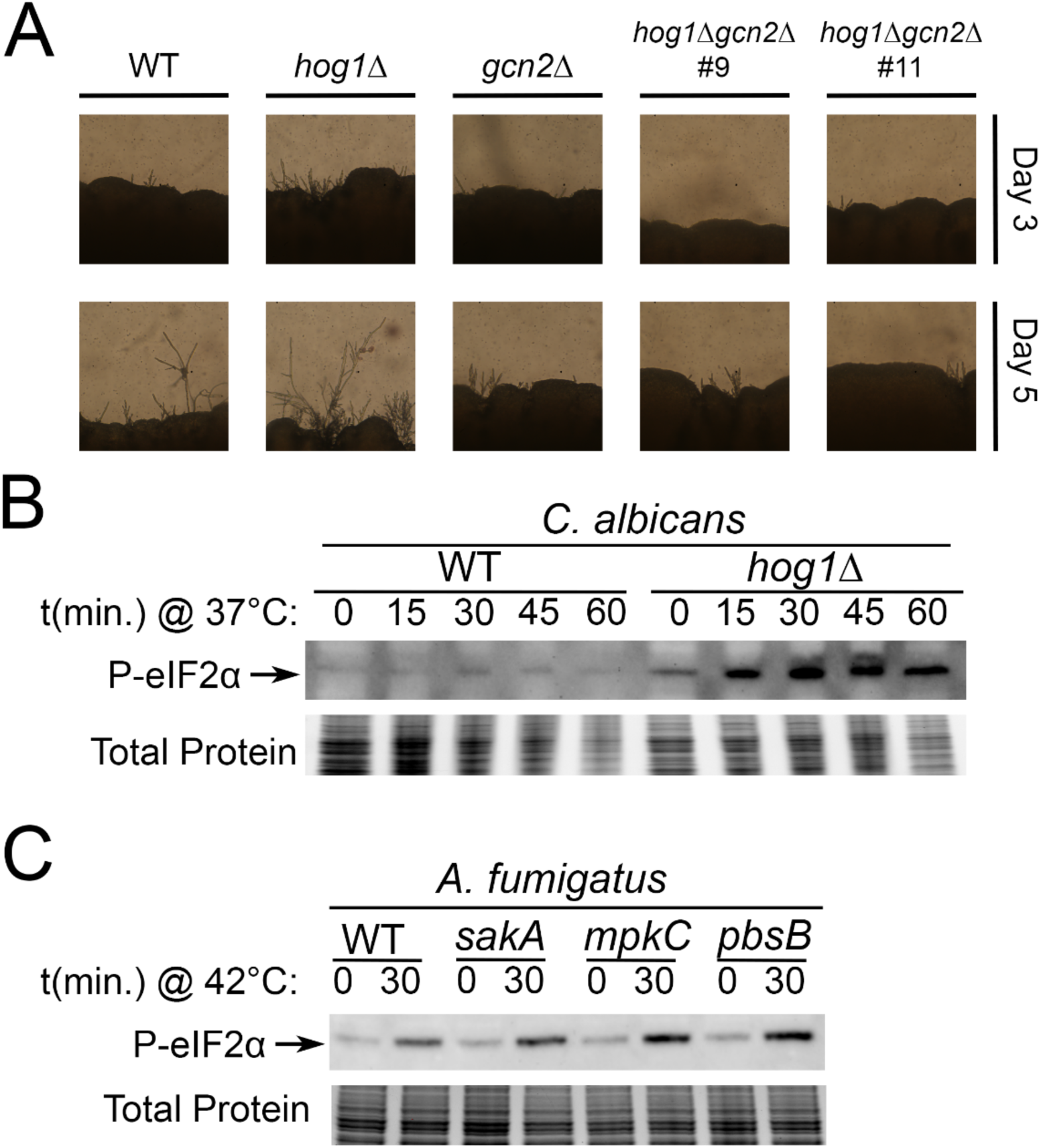
Conservation of Hog1-Gcn2 crosstalk in morphogenesis and other fungal pathogens. (A) The indicated MATα strains were mixed 1:1 with MATa KN99 *C. neoformans* and spotted on V8 agar and grown for 5 days. Representative photographs were taken after 3 and 5 days. (B) WT and *hog1*Δ DAY286 *C. albicans* were growth midlogarithmic phase in 30°C YPD media, followed by resuspension in 37°C YPD media. Protein samples were prepared and phosphorylated eIF2α was measured by western blotting, with total protein shown as a loading control. (C) WT, Δ*sakA,* Δ*mpkC,* and Δ*pbsB* CEA10 *A. fumigatus* were grown until germination in 37°C YPD media, followed by heat shock at 42°C for 30 minutes. Protein lysates were prepared and phosphorylated eIF2α was measured by western blotting, with total protein shown as a loading control.

To conclude our study, we wanted to address conservation of Hog1-Gcn2 crosstalk in other fungal pathogens. Although most other fungal pathogens of humans are ascomycetes, which diverged ∼400mya [15], there is evidence for crosstalk between these pathways in the model ascomycetes [26–28]. To test this, we measured P-eIF2α in WT and *hog1*Δ strains of *Candida albicans*, a pathogenic fungus belonging to the same order as *S. cerevisiae*. Relative to the parental WT strain, we observed greater P-eIF2α in the *hog1*Δ mutant during thermal stress (Fig. 7B). To further this exploration of ascomycetous fungal pathogens, we performed a similar analysis in *Aspergillus fumigatus*. The Hog1 pathway is bifurcated in *A. fumigatus*, with two Hog1 homologs, SakA and MpkC, acting downstream of a shared MAPK kinase, PbsB [29]. Thus, we used deletion mutants of both Hog1 homologs, as well as a PbsB-null strain to prevent activation of either kinase by the HOG pathway. We did not observe a substantial difference in the response of the WT background (CEA10) compared to *sakA*, *mpkC*, or *pbsB* (Fig. 7C). This data indicates that crosstalk between Hog1 and Gcn2 occurs in both *C. neoformans* and *C. albicans*, but not the obligate filamentous pathogen *A. fumigatus*. Considered as a whole, these results suggest crosstalk between Hog1 and Gcn2 regulates morphogenesis, and occurs in a species-specific manner over large evolutionary distances across the fungal kingdom.

## Discussion

Here we set out to investigate crosstalk and compensation between two major signaling modules, the Hog1 and Gcn2 pathways, which function concurrently during thermal stress. Our data demonstrate that crosstalk between these pathways regulates thermal stress adaptation, and improves our understanding of phenotypes associated with Hog1 deletion.

Considered as a whole, this study has important implications. Our data highlight the insufficiency of phenotypic characterization of deletion mutants to determine the function of genes. Several phenotypes associated with Hog1 deletion were, in fact, driven by differential regulation of the Gcn2 pathway rather than Hog1 itself. Thus, mechanistic studies are critical for understanding the etiology of an observed phenotype. This will allow us to develop more precise models of how genes function, and could reveal additional functions that are masked by compensatory activity. This is especially important to consider in the context of antifungal development, since this kind of information could improve predictions of which drug targets are likely to synergize. Additionally, studying compensation could identify pathways that often act as a fail-safe to conserve stress tolerance or virulence. This could vastly improve our understanding of how genes confer pathogenicity, as many could have been overlooked as virulence factors as a result of compensatory or redundant processes in knockout strains.

We first showed that loss of Hog1 was associated with increased phosphorylation of eIF2α and increased production of the ISR transcription factor, Gcn4 (Fig. 1A,C). Since we showed that phosphorylation of eIF2α is fully dependent on Gcn2 (Fig. 1B) and total eIF2α protein levels are not substantially altered in *hog1*Δ (Fig. S1), we propose two possible explanations: First, turnover of phosphorylated proteins, such as phospho-eIF2α or phospho-Gcn2, could be dysregulated in *hog1*Δ. This is plausible since there is evidence that Hog1 activity stimulates its own phosphatases in *S. cerevisiae* [30]. Second, there could be an increase in Gcn2 activity in *hog1*Δ. While this could conceivably result from increased levels of Gcn2 or its co-activators, loss of Hog1 could also increase levels of ribosome collisions or uncharged tRNAs, which activate Gcn2. One way this could occur is through uncontrolled translation elongation, which could increase collisions during stress or exhaust cellular pools of tRNAs. Indeed, Hog1 has been linked to repression of translation elongation via eEF2 phosphorylation in *S. cerevisiae* [13]. Alternatively, Hog1 could regulate processes upstream of Gcn2, such as tRNA charging or clearance of stalled ribosomes. Additional study of the precise mechanism underlying Hog1-Gcn2 crosstalk is warranted to improve our understanding of the nexus between these two pathways.

We associated our molecular observations with phenotypic analysis, which overall showed that double knockout of Hog1 and Gcn2 causes an exacerbated defect in kinetic growth during thermal stress compared to the single knockouts (Fig. 2). Notably, we did not observe a difference by spot plate analysis, and attribute the difference between these two approaches to the fact that spot plating is an end-point analysis, whereas growth curve analysis is kinetic.

In Figures 3-5, we revisited phenotypes that our lab previously reported in the *hog1*Δ mutant during thermal stress [12], and characterized the extent to which these phenotypes are dependent on the Gcn2 pathway. Collectively, these data suggest that Hog1-Gcn2 crosstalk is relevant to translation regulation and ribosomal transcript repression in *hog1*Δ, but is unimportant for cell wall and capsule phenotypes associated with Hog1 deletion. Thus, while Hog1 appears to regulate several classes of transcripts during thermoadaptation, only a subset of these exhibits a Gcn2-dependent contribution. We then moved to mechanistic questions about how Hog1 regulates signaling via the Gcn2 pathway (Fig. 6). Our results using fludioxonil and 3-AT suggest that Hog1 stimulation is sufficient to alter Gcn2 activation, and also indicates that crosstalk is not limited to the context of thermal stress. Additional research is warranted to determine whether Gcn2 rescues the response of Hog1 to other stressors that are relevant during infection.

In Figure 7, we briefly consider the broader implications of these findings in fungal biology. We found that Gcn2 was necessary for increased filamentation of *hog1*Δ (Fig. 7A). Given the well-established role of Hog1 as a regulator of growth morphology in several fungi [16,24,25], we suspect that Gcn2 may contribute to morphology of Hog1-deleted strains in other fungi. Indeed, a previous study in *C. albicans* found the ISR promotes filamentation during amino acid starvation [31].

Given the conserved role of Hog1 in fungal virulence, we sought to determine whether Hog1-Gcn2 crosstalk is also conserved in other fungal pathogens, by assessing eIF2α phosphorylation in *hog1*Δ mutants in *C. albicans* and *A. fumigatus* (Fig. 7B-C). We found that Hog1 deletion upregulated Gcn2 signaling in *C. albicans*, but did not have a substantial effect in *A. fumigatus*. Our findings in *C. albicans*, an ascomycete fungus, suggest that linkage between the Hog1 and Gcn2 pathways can be maintained over long evolutionary distances (∼400 million years). This is also supported by previous work in the model fungi *S. cerevisiae* and *S. pombe*, which have increased P-eIF2α in response to stress when the Hog1 homolog is deleted [26–28]. While the response we observed in *A. fumigatus* appears incongruent with our findings and previous studies, we suspect this due to differences in the underlying biology. Among other things, *A. fumigatus* has an obligate filamentous growth program and is adjusted to growth at temperatures far exceeding mammalian core temperature. Thus, it seems unsurprising that signaling would proceed differently in this species.

Overall, this study advances our understanding of the Hog1/p38 pathway, and points to important considerations for the study of core conserved signal transduction pathways. Given the broad specificity of major pathways that emerged early in the eukaryotic lineage (e.g. TOR, PKA, GCN, MAPKs), it’s critical that we consider how they impact one another, and characterize the biological basis of their interactions. Understanding the cause-and-effect relationships between pathways will improve our models of stress responses, as well as our ability to manipulate these processes as therapeutic targets.

## Materials and Methods

### Strains

*C. neoformans* strain H99 was used as the wild-type background. The *hog1*Δ mutant was constructed using TRACE [32]. Plasmids pYF24 and pDD162 were obtained from the Lin Lab at the University of Georgia. Briefly, the *hog1*Δ knock-out construct was amplified from gDNA of a *hog1*Δ mutant in a related parental background (Joe Heitman, Duke University) using primers F-HOG1KOamp (TGACTTTGGCGCTTGTTGC) and R-HOG1KOamp (GCAGATTGACGAATTCACCC). The gRNA construct was amplified by TRACE protocol using *HOG1*-specific primers R-HOG1g+U6 (agctgctcggtactgaccttAACAGTATACCCTGCCGGTG) and F-HOG1g+sgTERM (aaggtcagtaccgagcagctGTTTTAGAGCTACAAATAGCAAGTT). Electroporation was performed with 100ng Cas9 cassette, 100ng sgRNA cassette, and 2ug knockout construct. Recombination was verified by PCR, and northern blot confirmed ablation of *HOG1* expression. This approach was used with biolistic transformation to generate *hog1*Δ*gcn2*Δ mutants, which were made using a previously described G418-resistant *gcn2*Δ as a parental strain [10]. *pbs2*Δ, *ssk2*Δ, and *mpk1*Δ strains were obtained from the Bahn kinase deletion collection [33], and *ssk1*Δ from the Madhani deletion collection via the Fungal Genetics Stock Center (FGSC) [34]. *C. albicans hog1*Δ and the parental WT were obtained from Aaron Mitchell’s kinase deletion collection at the FGSC [35]. *A. fumigatus* Δ*sakA,* Δ*mpkC,* Δ*pbsB*, and the parental WT CEA10 strain were obtained from the *A. fumigatus* kinase deletion collection (Jarrod Fortwendel, University of Tennessee Health Science Center) [36].

### Media and growth conditions

Unless otherwise stated, cells were grown overnight in 30°C YPD (1% yeast extract, 2% peptone, and 2% dextrose) with 250rpm shaking in snap-cap tubes. For growth curves, spot plates, capsule, and mating studies, cells were used directly from overnight cultures. For western and northern blotting, qPCR, and polysome profiling, cells were seeded at OD_600_ = 0.2 and grown in flasks to midlogarithmic phase (OD_600_ ∼ 0.6) in 30°C YPD at 250rpm prior to treatment. Stress at 37°C was introduced by pelleting and resuspension in fresh pre-warmed YPD and incubated with shaking in a 37°C incubator. For spot plates, cells were plated on YPD agar (YPD, 2% agar) with the indicated drugs and incubated at the indicated temperatures. For mating, cells were plated on V8 agar (5% V8 juice, 4% agar, 0.5g/L KH_2_PO_4_) and incubated in the dark at 21°C surrounded by CaSO_4_ dessicant. For capsule cells were grown overnight in 0.1x Saboraud dextrose media (10% Sabouraud pH 7.3, 50mM MOPS) at 37°C in snap-cap tubes with shaking.

### Growth analyses

For spot plates, cells were washed with sterile deionized water, followed by resuspension at OD_600_ = 1.0. A ten-fold dilution series was performed, and 5μL of each dilution was spotted. For temperature sensitivity analysis, cells were plated on YPD agar and incubated at 30°C, 37°C, or 38°C. For cell wall stress analysis, cells were plated on 0.5mg/mL caffeine, 1mg/mL calcofluor white, 5mg/mL Congo red, 0.15%-0.3% sodium dodecyl sulfate (SDS), or no drug controls. Plates were photographed after three days and one of three biological replicates is shown.

For growth curves, cells were suspended at OD_600_ = 0.1 in fresh YPD. 100μL of cells were plated in triplicate wells of a 96-well plate and grown in a Bio-Tek Synergy Neo plate reader with linear shaking (567cpm) at the indicated temperatures. The OD_600_ was read every hour for 24hrs. Growth curves show a representative replicate. Growth rate during exponential growth (∼8-13h) was calculated across three biological replicates as follows: Rate = ln(OD_600_(final)/ OD_600_(initial))/duration.

### Western blotting

15mL of cells were collected at the indicated time points, pelleted, and flash-frozen. Thawed cells were lysed and SDS-PAGE was performed as described previously [12]. P-eIF2α was probed using rabbit α-phospho-eIF2α (AbCam, ab32157) and an HRP-conjugated secondary antibody (CST, #7074). Total eIF2α and Gcn4 levels were detected with custom-generated antibodies (GenScript) raised in rabbits exposed to full-length recombinant protein. HRP signal was measured on a BioRad ChemiDoc MP using Clarity Max ECL substrate (Bio-Rad). Each image is representative of three biological replicates. Densitometry was performed using ImageLab software (Bio-Rad).

### qPCR

5mL of cells were collected after 1hr stress at 37°C and flash-frozen. Cells were lysed and RNA isolation and cDNA synthesis was performed as described previously [12]. qPCR for *CHS4* (CNAG_00546), *CHS5* (CNAG_05818), *EXG1* (CNAG_05803), and mitofusin (CNAG_06688) was performed using previously reported primers [12] using the qPCR SyGreen Blue mix LoROX (PCR Biosystems). Signal was measured using a CFX Connect real-time PCR detection system (Bio-Rad).

For each experiment, two technical replicates were measured, and the average C_T_ value was used for quantification. Mitofusin served as an internal control, and ΔΔC_T_ was used to calculate relative expression. Bar graph depicts the average of three biological replicates.

### Capsule staining

Strains grown in capsule-inducing media were mixed 1:5 with India ink and then subjected to differential interference contrast (DIC) microscopy. 50 cells per replicate were imaged with a 63x DIC objective on a Leica DMi 8 inverted microscope. Measurements of capsule size were performed using the freehand line and measurement tools in ImageJ. Bar graph depicts capsule thickness of fifty cells from one of three biological replicates.

### Polysome profiling

Cells were left untreated or grown for 30 minutes at 37°C, pelleted, and flash-frozen. Cells were lysed by mechanical disruption in polysome lysis buffer (20 mM Tris-HCl pH 8.0, 140 mM KCl, 5 mM MgCl_2_, 1% Triton X-100, 25 mg/mL heparin sodium sulfate, 0.1 mg/mL cycloheximide) and RNA concentration in lysate was measured on a NanoDrop One. 200μg RNA was loaded over a 10-40% sucrose gradient, followed by centrifugation at 39,000rpm for 2 hrs in a SW41-Ti swinging bucket rotor. The gradient was then run through the Piston Gradient Fractionator (Biocomp), which measured absorbance at 260nm. The graphs show one of three biological replicates. The polysome-to-monosome was calculated by manually defining the boundaries of these regions of each profile and calculating the area-under-curve using Prism software. These ratios were normalized the WT sample for each condition.

### Northern blotting

5mL cells were pelleted and flash-frozen at the indicated time points. RNA was harvested and the *RPL2* transcript was probed by Northern blot using a ^32^P-labeled probe as described previously [12]. Signal from the blot was obtained using a phosphor screen with a Typhoon imager (GE). This signal was quantified by densitometry using ImageLab (BioRad) and normalized to rRNA signal. Blots show one representative replicate, and bar graph shows signal after one hour of thermal stress, normalized to start, across three biological replicates.

### Filamentation

Cells from overnight cultures were set to an OD_600_ = 1.0 in distilled water and mixed 1:1 with MATa KN99. 20μL of the resulting mixture was spotted on V8 agar and incubated in dark, dry conditions at 21°C for 5 days. Filamentation at the colony periphery of the spot was photographed on days 3 and 5 using a Sony DSLR-A300 camera attached to an Olympus BX41 microscope at 40X magnification. Data shown is representative of two biological replicates where three mating spots were photographed.

### Candida and Aspergillus experiments

For *Candida* experiments, cells and samples were treated exactly as described for *Cryptococcus*, with the exception that during lysis a shorter duration of bullet blending was needed. For *Aspergillus*, conidia were harvested from plates and stored overnight at 4°C in sterile water. The next day, conidia were added to YPD media and incubated at 37°C with shaking until germ tube formation (∼8 hrs), after which the cultures were subjected to heat shock by resuspension in 42°C media for thirty minutes. Protein lysates were prepared as described for *Cryptococcus*.

### Statistical analysis

Bar graphs were made in Prism software with error bars representing the standard deviation (SD). All statistical tests were performed by one-way ANOVA with Tukey’s test *post hoc,* with significance cutoffs were defined as *\*, P* < 0.05; *\*\**, *P* < 0.01; *\*\*\**, *P* < 0.001; ****, *P* < 0.0001.

## Acknowledgements

1. *C. neoformans* and *C. albicans* strains were obtained from the Fungal Genetics Stock Center (Manhattan, KS, USA). The *A. fumigatus* strains were a gift from Jarrod Fortwendel (University of Tennessee, Health Science Center). This work was funded by Insititutes of Allergy and Infectious Disease at the National Institute of Health under award numbers R01AI131977 and F31AI169969.

---

**Fig S1:**
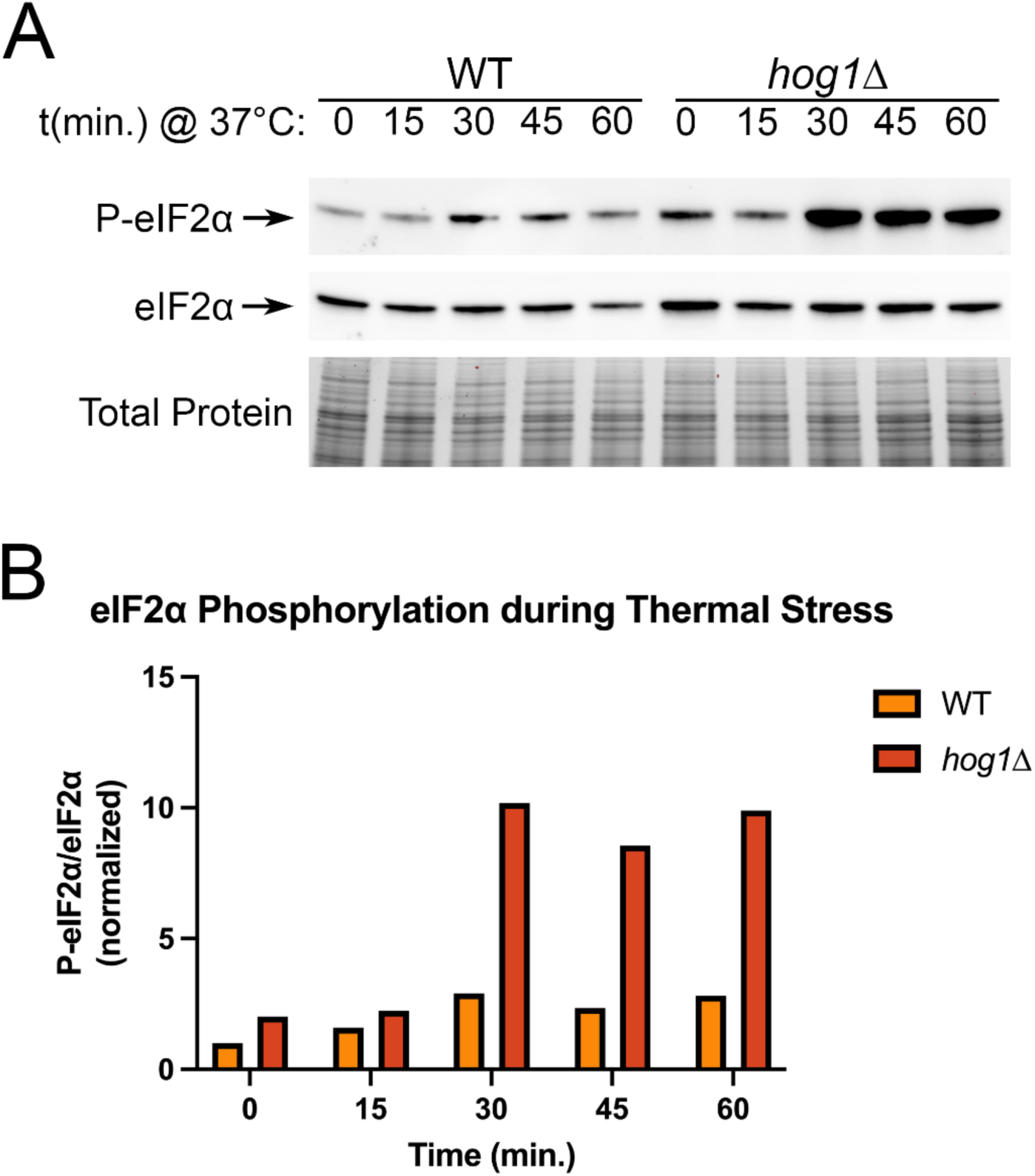
A greater proportion of eIF2α is phosphorylated in *hog1*Δ during thermal stress. (A) The indicated strains were grown to midlogarithmic phase at 30°C in YPD media, followed by resuspension in pre-warmed 37°C YPD media. Samples were collected at the indicated time points, and protein samples were prepared and probed by western blotting using an antibody against total eIF2α or phosphorylated eIF2α. Total protein is shown as a loading control (*n = 3*). (B) Quantification of signal from panel A, with phosphorylated eIF2α normalized to total eIF2α.

**Fig S2:**
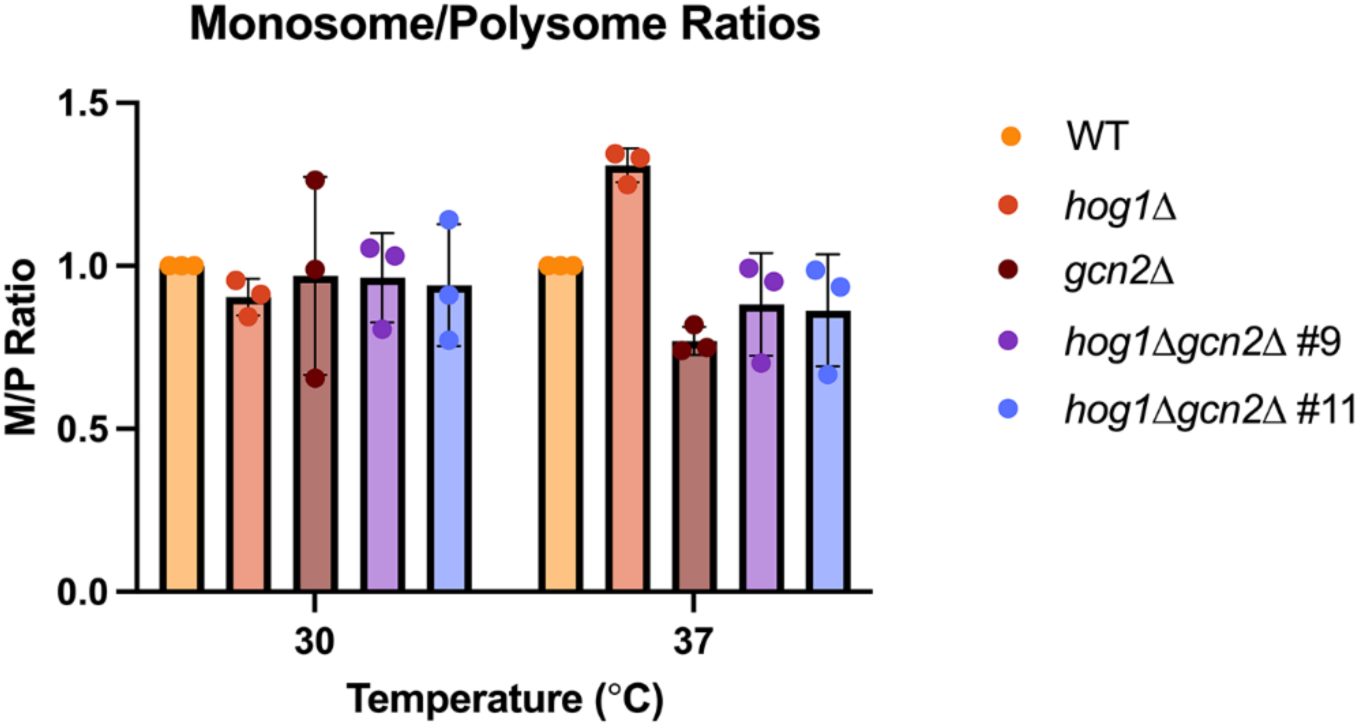
Quantification of polysome profiles during thermal stress. Polysome profiling was performed as described in Figure 4. The monosome-to-polysome (M/P) ratio was calculated by manually defining the boundaries of these sections of the polysome profile and performing AUC analysis using Prism software. The results for three biological replicates are shown (*n = 3*). Error bars show standard deviation.

